## Supplementary Figures for "Modular actin nano-architecture enables podosome protrusion and mechanosensing"

Van den Dries et al.

### Van den Dries et al. Supplementary Figure 1

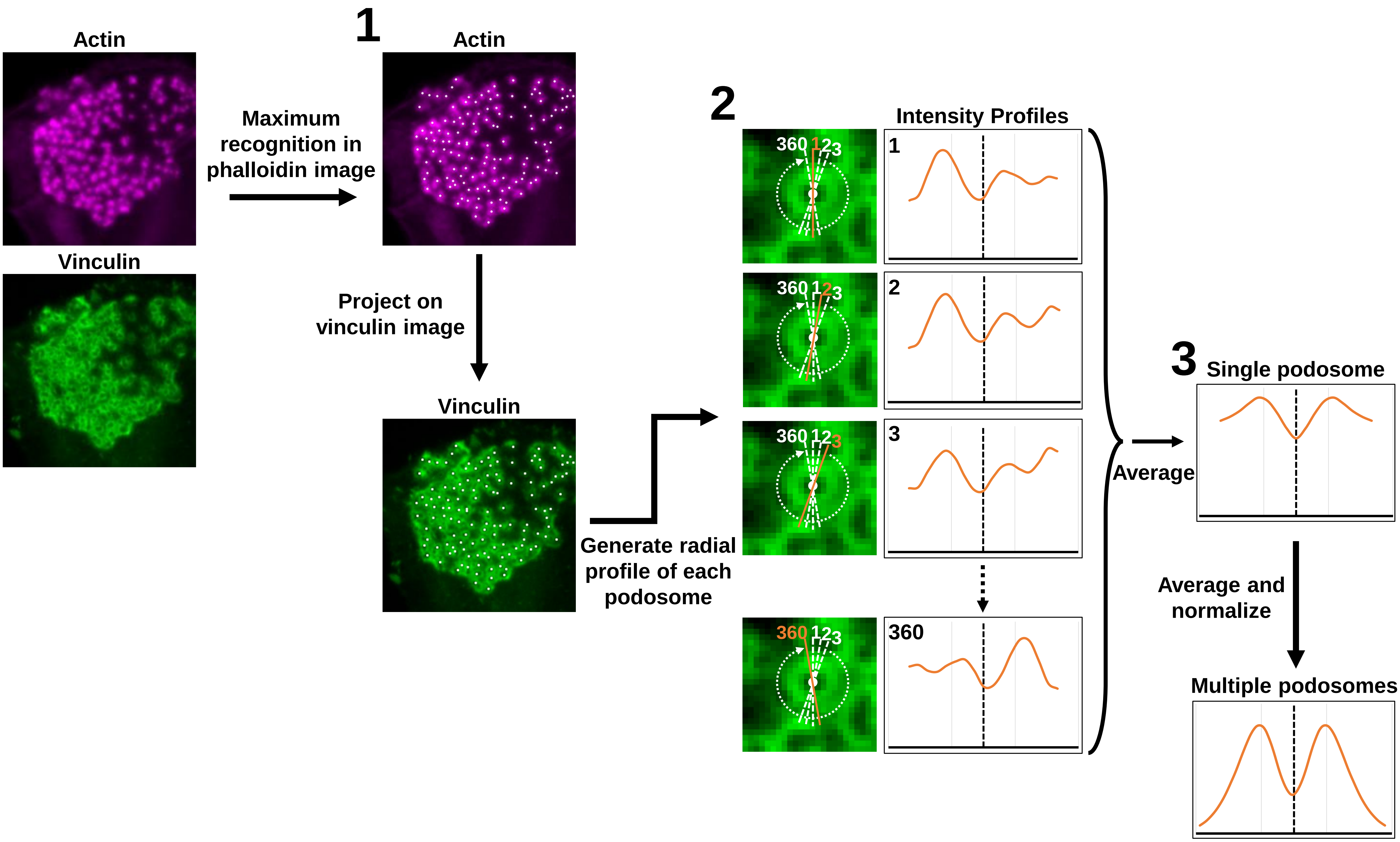


**Supplementary Figure 1**

**Overview of the radial fluorescence profile analysis**

To quantify the localization of each of these proteins with respect to the podosome core we used a semi-automatic self-developed ImageJ macro that **1**) recognizes the podosome core centers based on the actin image **2**) draws a vertical line of ~3um through the center of the core that rotates around its center and collects a profile for every line and **3**) produces an average radial intensity profile as a function of distance from the podosome core center. Profiles are normalized to the minimum and maximum for visualization and comparison. Of note, for some of the panels we only present one half of the intensity profile since the radial intensity profiles are symmetric by definition.

### Van den Dries et al. Supplementary Figure 2

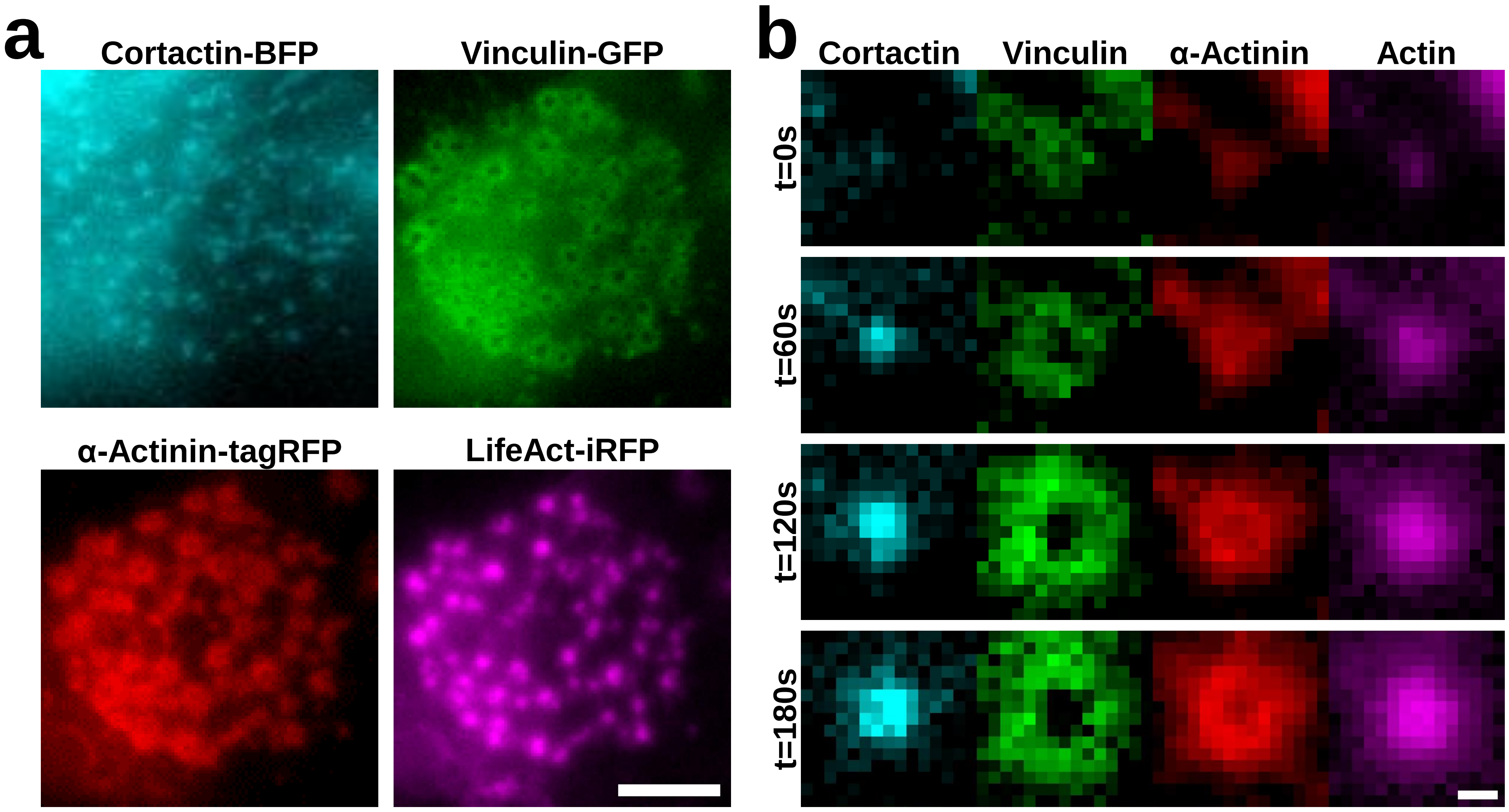


**Supplementary Figure 2**

**Simultaneous visualization of different actin-binding proteins at podosomes in living cells**
**a-b,** Widefield images of a DC transfected with cortactin-BFP (cyan), vinculin-GFP (green), α-actinin-tagRFP (red) and LifeAct-iRFP (magenta). **a,** shows a still from an entire podosome cluster and **b,** shows a few stills in time of a single assembling podosome. Scale bars: **a** = 5 µm, **b** = 0.5 µm

### Van den Dries et al. Supplementary Figure 3

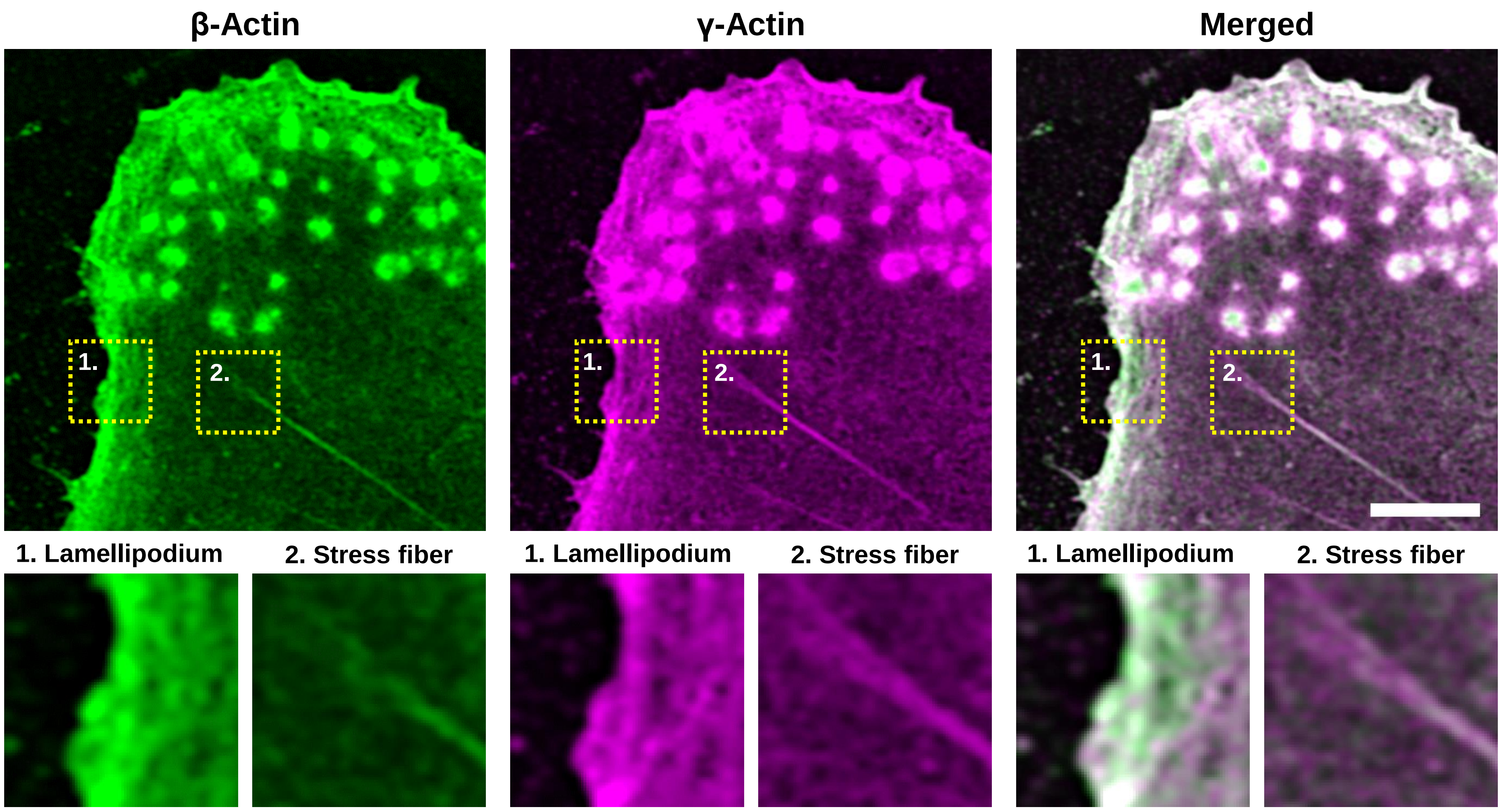


**Supplementary Figure 3**

**Distribution of γ and β-actin in whole cells**

**a,** Airyscan images of a DC stained for γ (magenta) and β-actin (green). Insets in the left corner depict a lamellipod that is enriched in β-actin and insets in the right corner depict a stress fiber that is enriched in γ-actin. Scale bar = 5 µm.

### Van den Dries et al. Supplementary Figure 4

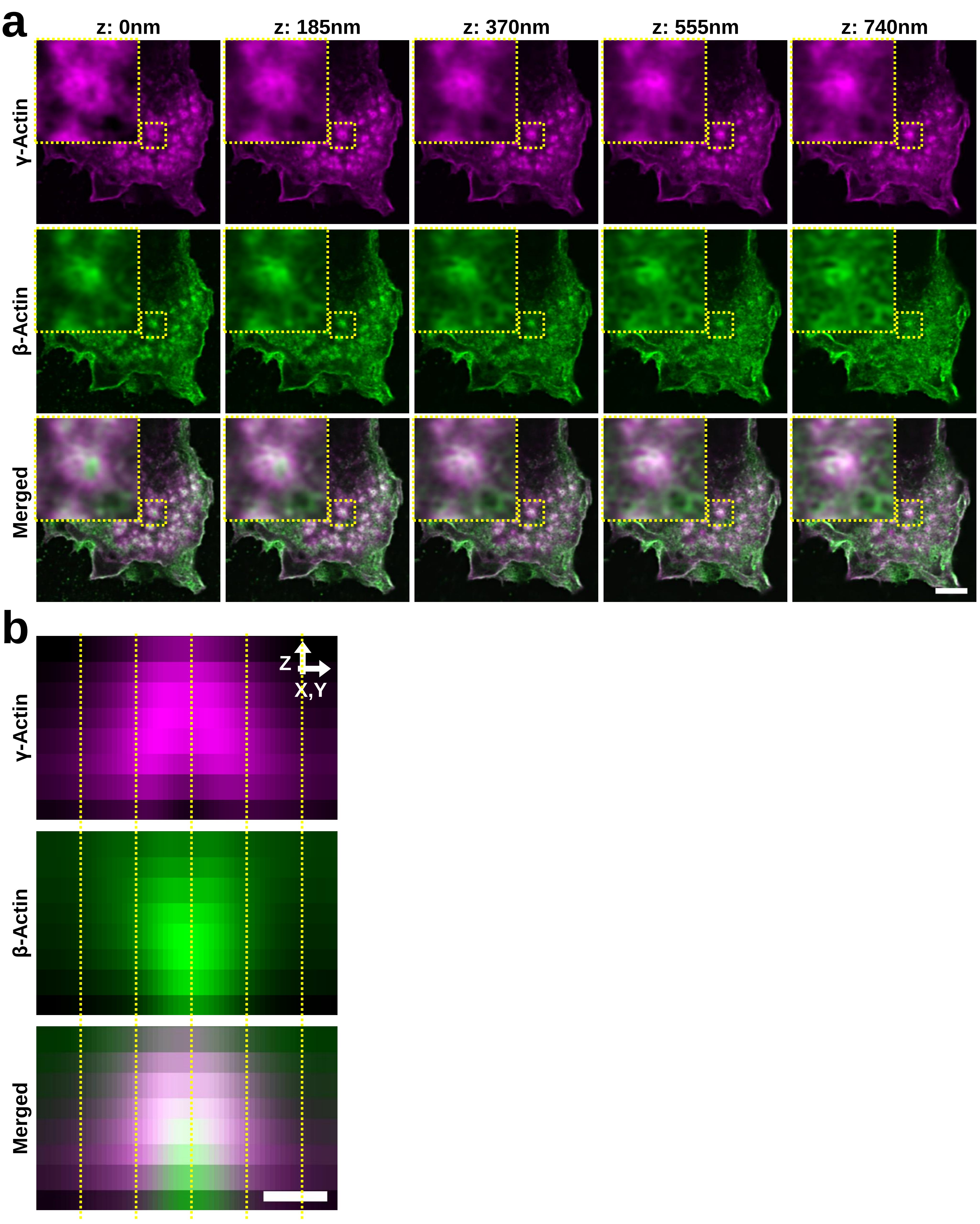


**Supplementary Figure 4**

**γ and β-actin isoforms differentially localize to cPM and pPM within podosomes of murine BMDCs**
**a,** 3D-Airyscan images of a murine BMDC stained for γ (magenta) and β-actin (green). Insets depict a single podosome. **b,** Average radial orthogonal view of γ (magenta) and β-actin (green) in 10 podosomes. Bottom panel shows the merged image. Scale bars: **a** = 2 µm, **b** = 0.5 µm.

### Van den Dries et al. Supplementary Figure 5

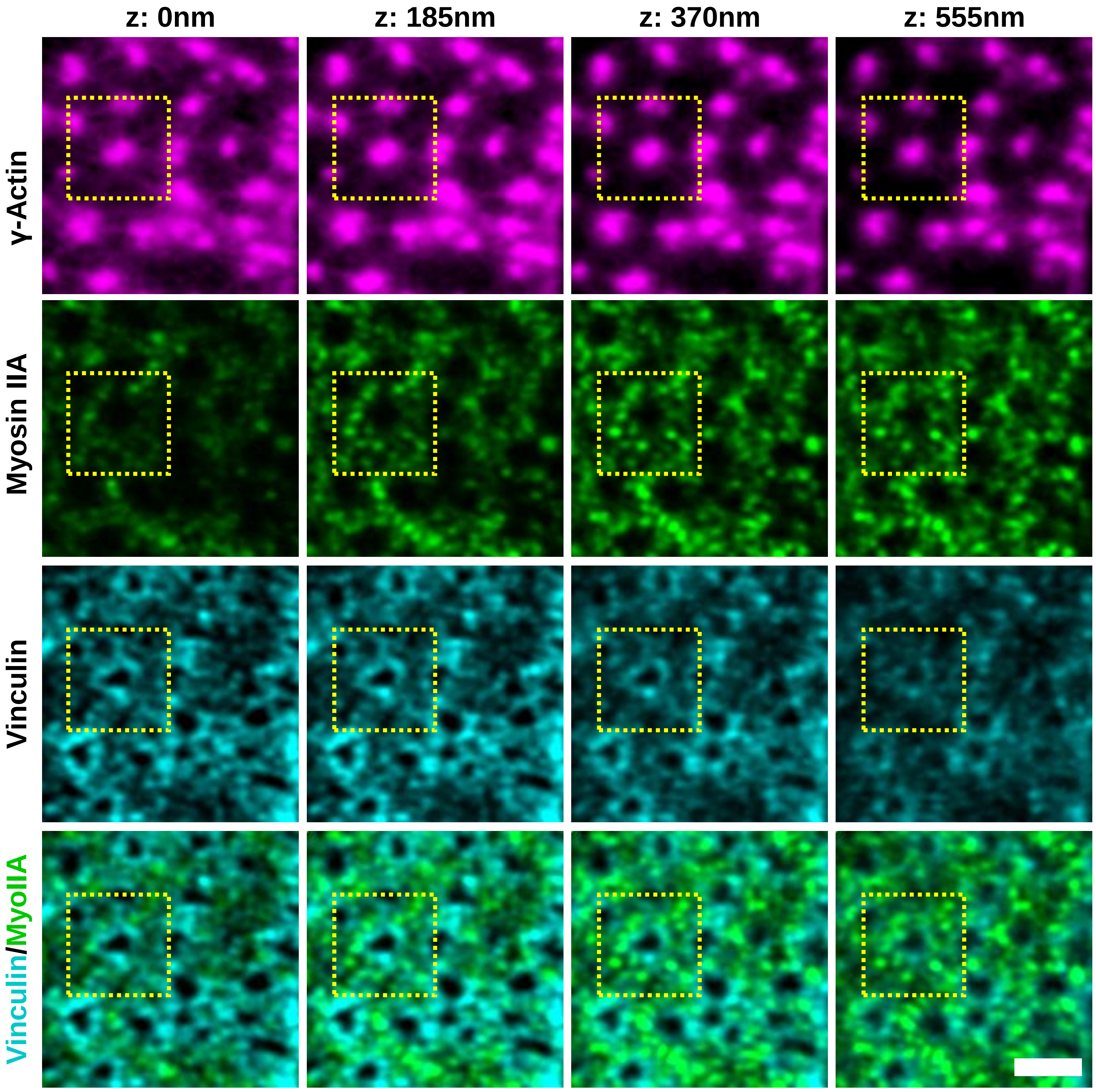


**Supplementary Figure 5**

**Vinculin and myosin IIA occupy distinct regions in podosome clusters**

3D-Airyscan images of a DC stained for actin (magenta), myosin IIA (green) and vinculin (cyan). Images shown are the total clusters for the single podosome shown in main Figure 3a. Podosome shown in main Figure 3a is indicated by a the yellow dashed square. Scale bar = 2 µm.

### Van den Dries et al. Supplementary Figure 6

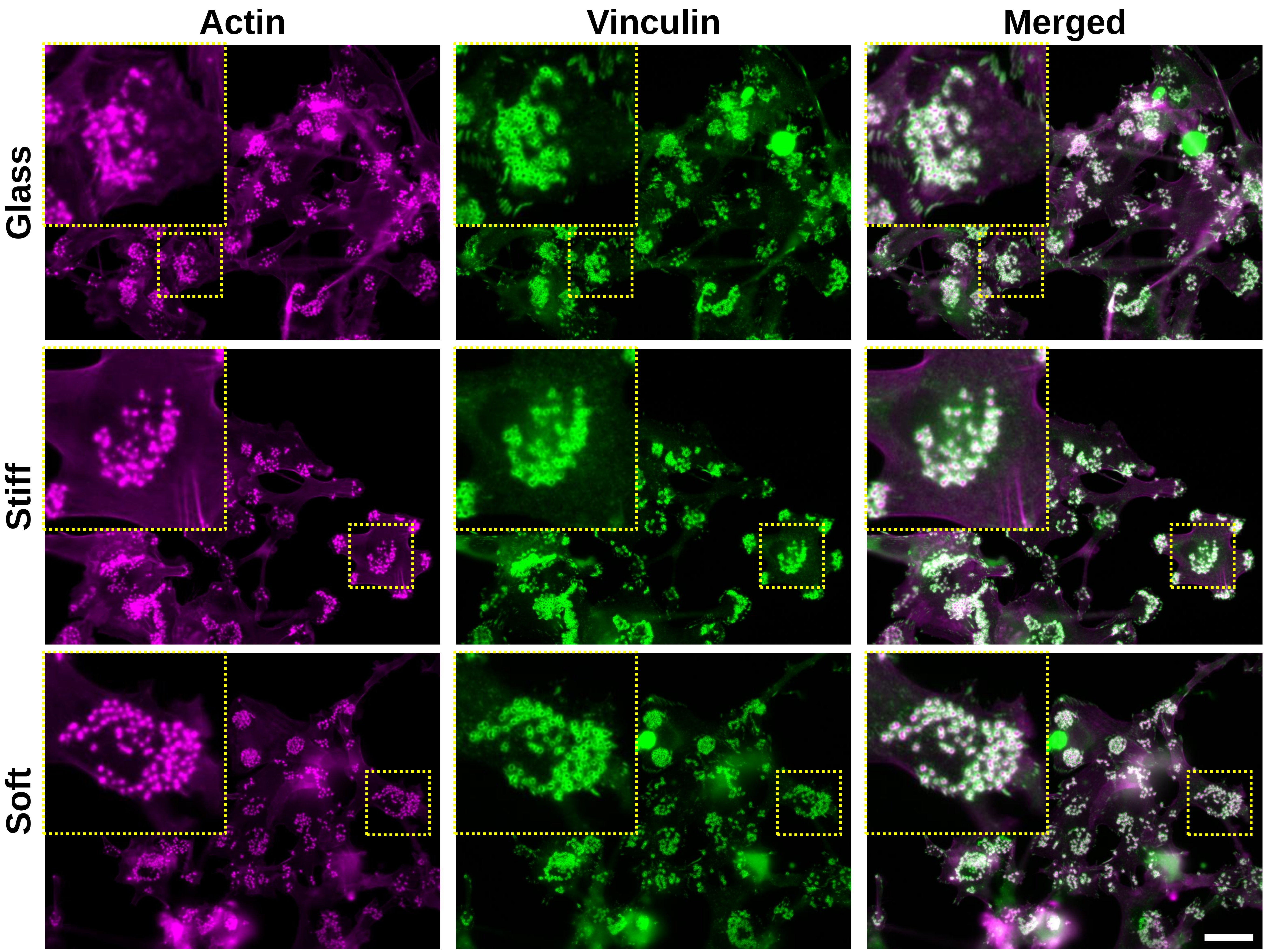


**Supplementary Figure 6**

**DCs spread and form podosomes independent of substrate stiffness**
DCs were seeded on glass, stiff and soft PDMS and fixed after 3 hrs. Shown are widefield images of DCs stained for actin (magenta) and vinculin (green). Insets depict single cells and the right panel depicts the merged image. Scalebar = 25 µm.

### Van den Dries et al. Supplementary Figure 7

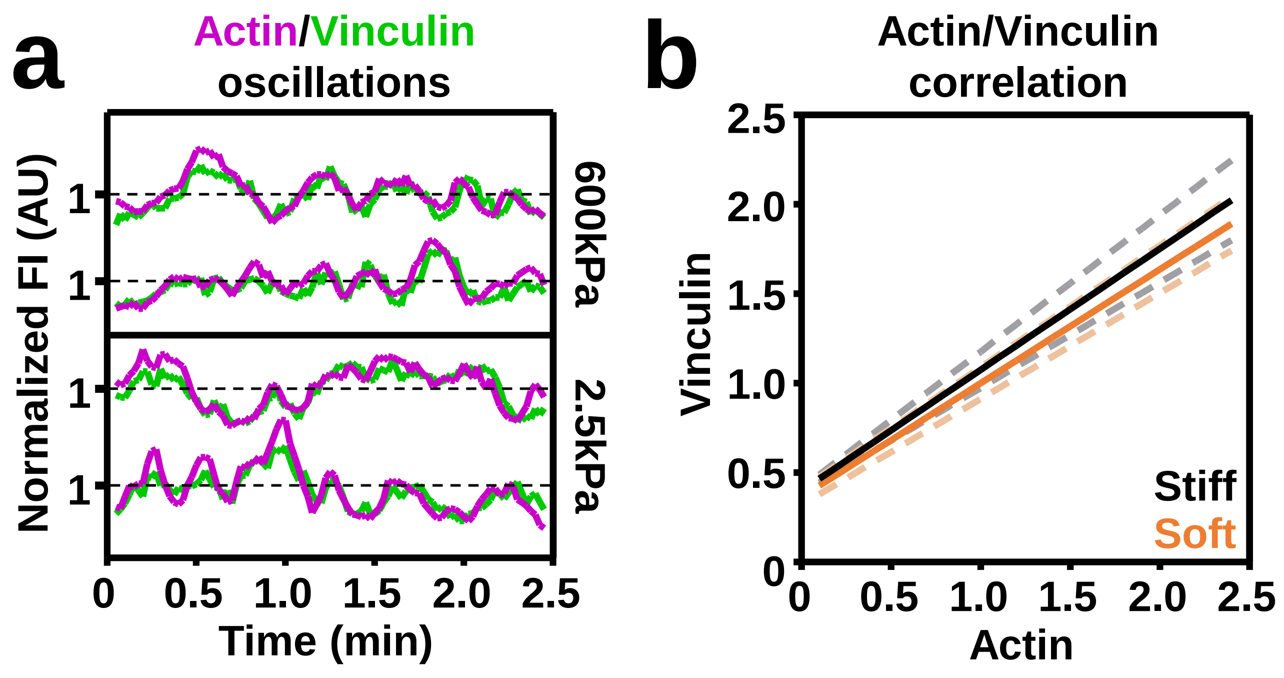


**Supplementary Figure 7**

**Podosome protrusive forces only deform soft PDMS**
**a,** DCs were transfected with Lifeact-GFP and vinculin-mCherry. Imaging was performed with Airyscan confocal microscopy with 15s frame intervals. The graph depicts oscillations of the fluorescence intensity of Lifeact-GFP (magenta) and vinculin-mCherry (green) in representative podosomes on stiff (upper panel) and soft (lower panel) substrate over time. **b,** Lifeact-GFP and vinculin-mCherry fluorescence intensity was correlated and fitted using a linear regression analysis on both stiff ($r^2$=0.51, P<0.0001) and soft ($r^2$=0.77, P<0.0001) substrates.

### Van den Dries et al. Supplementary Figure 8

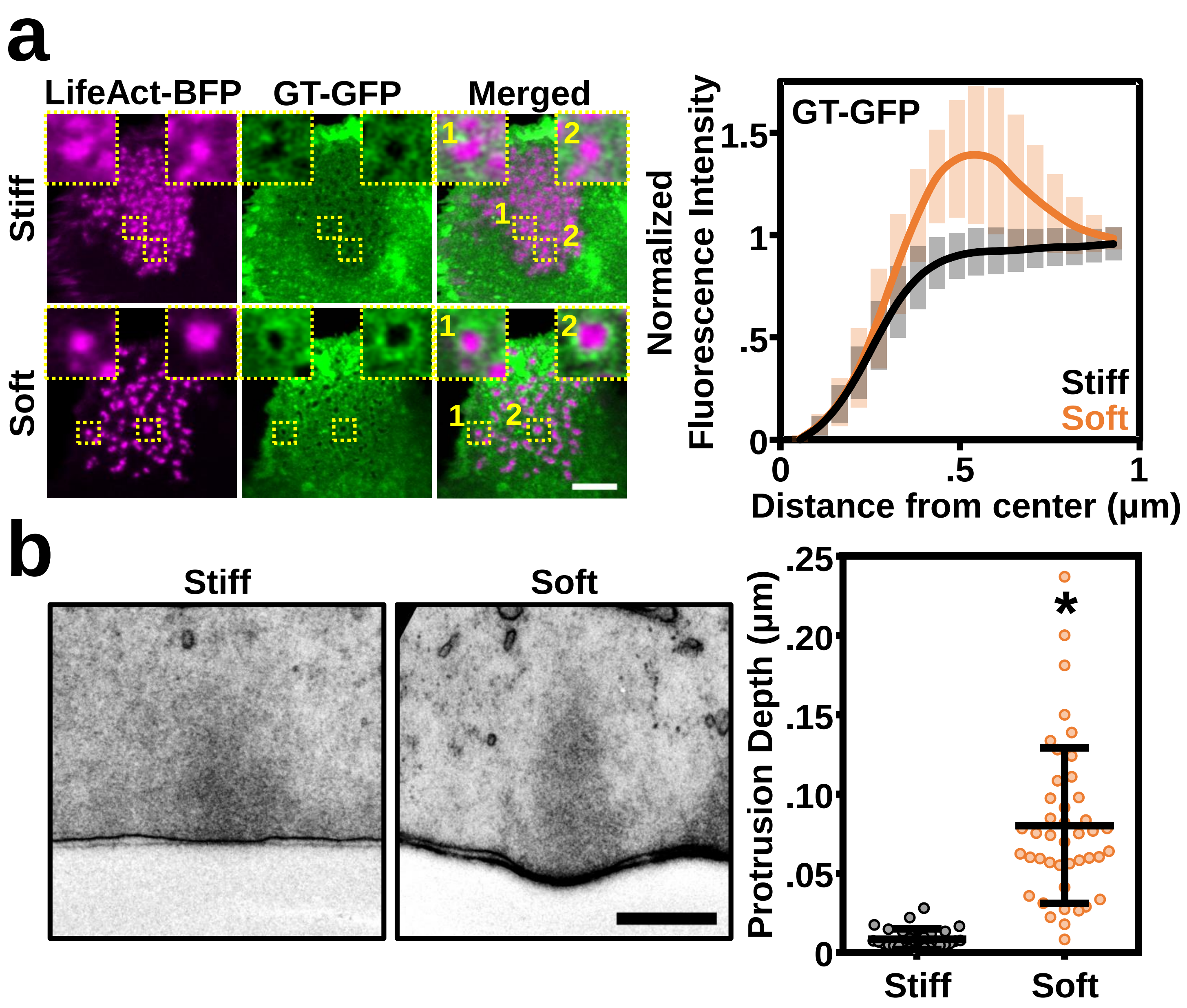


**Supplementary Figure 8**

**Podosome protrusive forces only deform soft PDMS**

**a,** Airyscan images of DCs transfected with LifeAct-BFP and GT-GFP. The graph shows the average radial fluorescent intensity profile of GT-GFP on stiff (black) and soft (orange) substrates. **b,** Transverse EM sections of single podosomes showing no protrusion on the stiff substrate and a small protrusion on the soft substrate. The graph depicts a quantification of the protrusion depth on both stiff and soft substrates. Scale bars: **a** = 5 µm, **b** = 0.5 µm.

### Van den Dries et al. Supplementary Figure 9

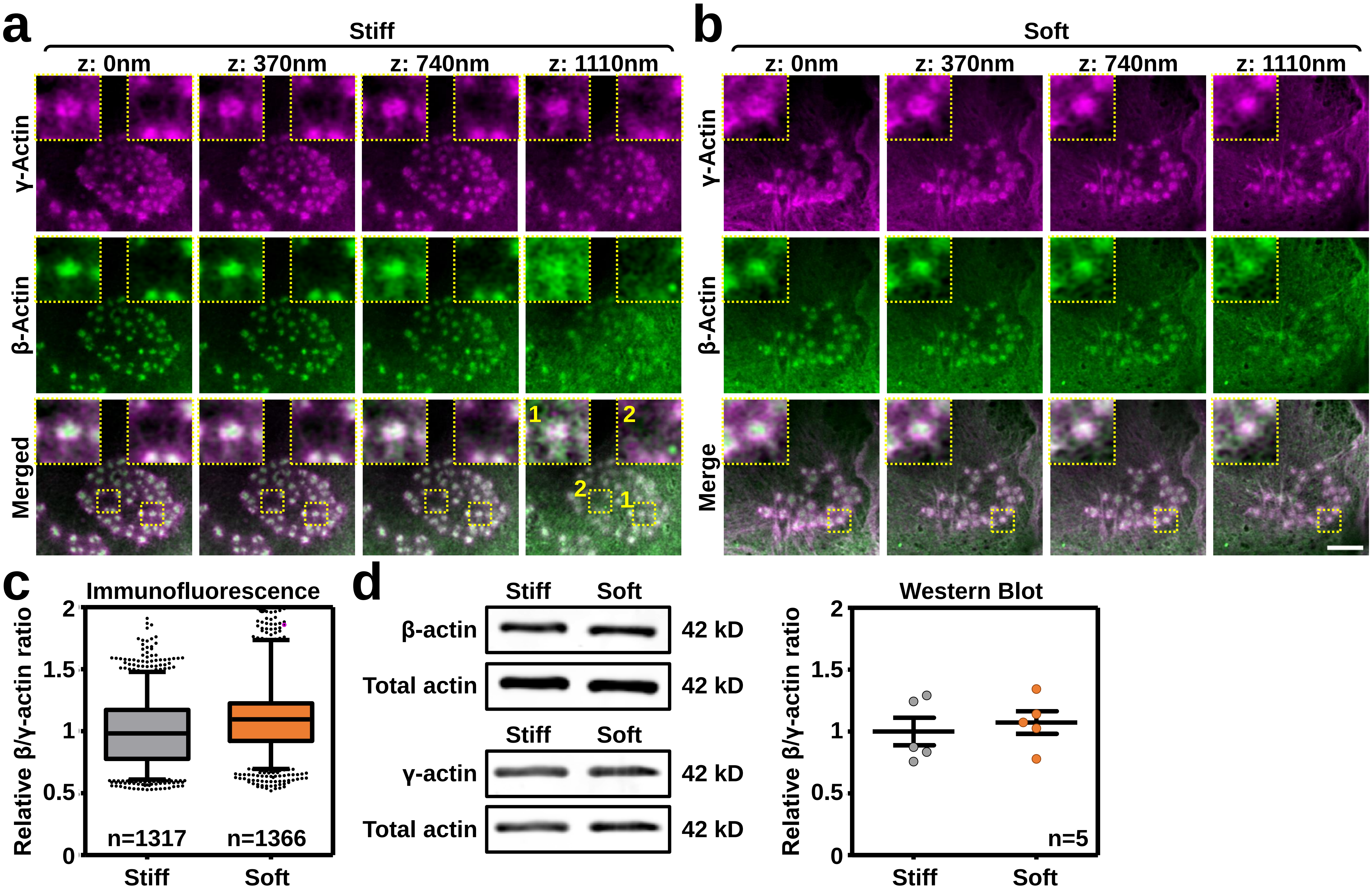


**Supplementary Figure 9**

**Differential localization and relative levels of β and γ-actin are independent of substrate stiffness**
**a-b,** 3D-Airyscan images of DCs seeded on **a**, stiff and **b**, soft substrates and stained for γ (magenta) and β-actin (green). Insets depict a single podosome. Note the localization of γ-actin to both the pPM (inset 1) and the radiating filaments (inset 2) on stiff substrates. Scale bar = 5 µm. **c,** Quantification of the β/γ actin ratio on soft (n = 1317 podosomes) and stiff (n = 1366 podosomes) substrates. **d,** Shown is a western blot analysis of β and γ-actin of lysates prepared from VPMs of DCs on stiff and soft substrates. Total actin is used as a loading control. The graph depicts a quantification of the β/γ actin ratio in 5 independent experiments.

### Van den Dries et al. Supplementary Figure 10

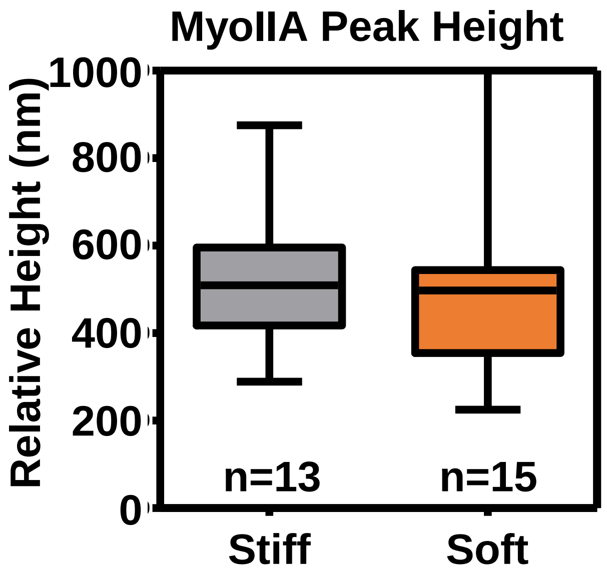


**Supplementary Figure 10**

**Myosin IIA peak intensity not controlled by substrate stiffness.**
Quantification of the fluorescence peak height of myosin IIA on soft (n=13 clusters) and stiff (n = 15 clusters) substrates.

### Van den Dries et al. Supplementary Figure 11

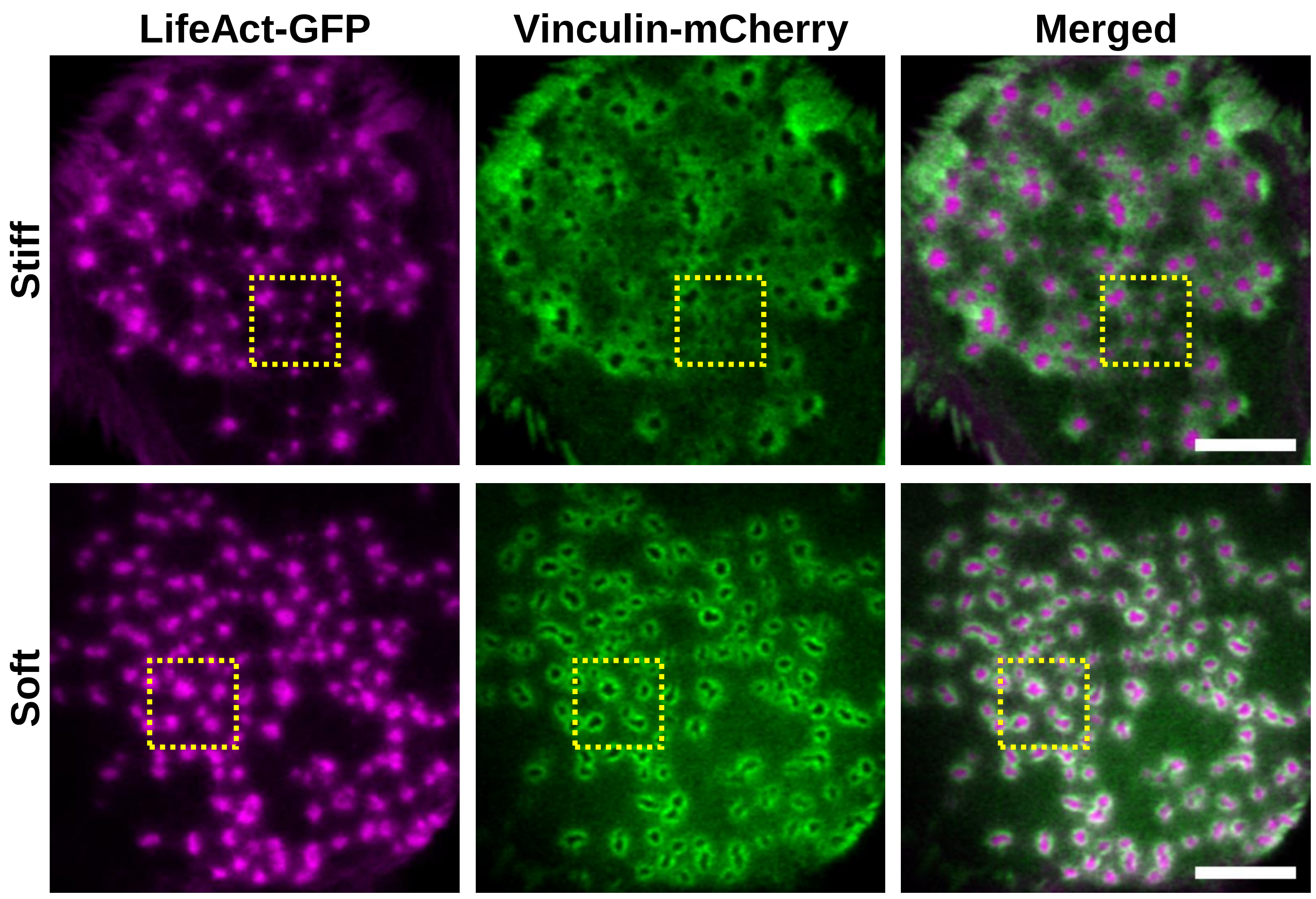


**Supplementary Figure 11**

**Actin and vinculin reorganize on soft substrates in living cells**
Airyscan images of DCs transfected with Lifeact-GFP (magenta) and Vinculin-mCherry (green) and seeded on stiff and soft substrates. The yellow dashed square indicates the area depicted in main Figure 6c. Scale bar = 5 µm.

### Van den Dries et al. Supplementary Figure 12

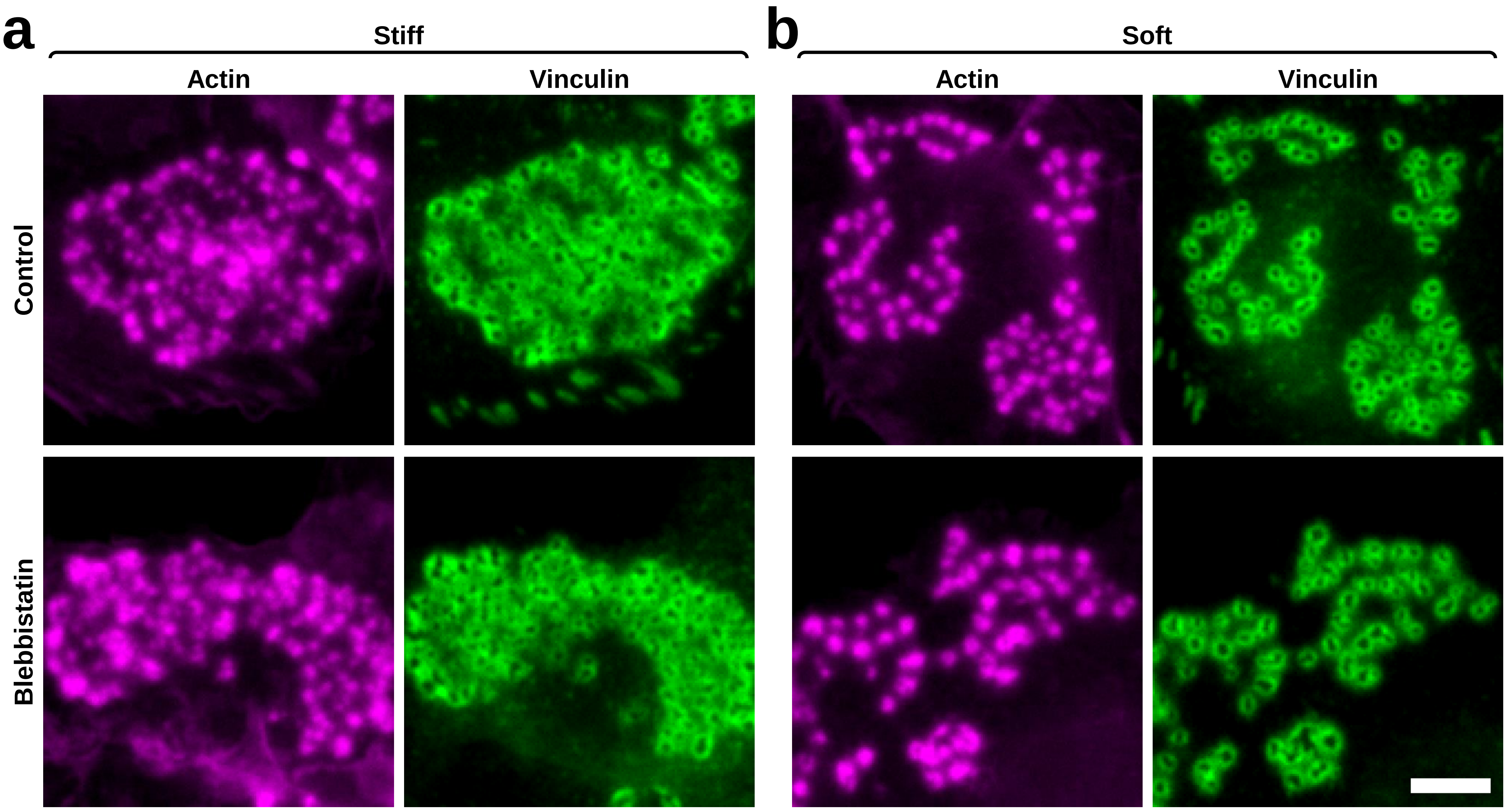


**Supplementary Figure 12**

**Myosin IIA does not control the stiffness induced relocalization of vinculin**
Widefield images of DCs seeded on stiff and soft substrates, fixed and stained for actin (magenta) and vinculin (green). Cells were either left unstimulated (top panels) or treated with 50 μM Blebbistatin for 1 hr (bottom panels) prior to fixation. Scale bar = 5 μm.

### Van den Dries et al. Supplementary Figure 13

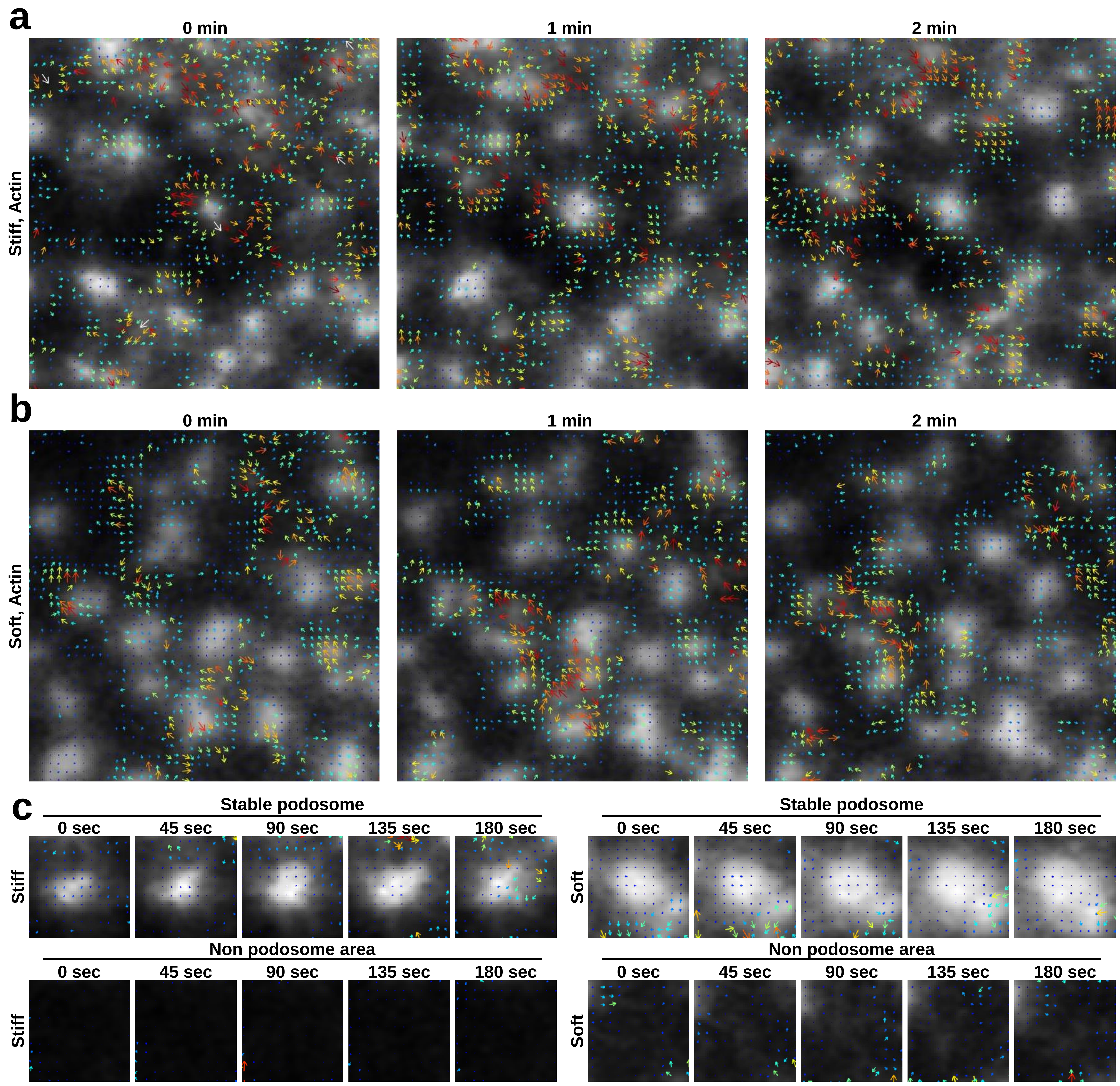


**Supplementary Figure 13**

**twSTICS reveals flows of actin in dynamic podosomes on stiff and soft substrates**

**a-c,** DCs were transfected with Lifeact-GFP and vinculin-mCherry. Airyscan imaging was performed with 15s frame intervals. Time series were subjected to twSTICS analysis and results are plotted as vector maps in which the arrows indicate direction of flow and both the size and color are representative of the flow magnitude. Shown are representative waves of vectors for actin on **a,** stiff and **b,** soft substrates as well as in **c,** stable and non podosome areas.

### Van den Dries et al. Supplementary Figure 14

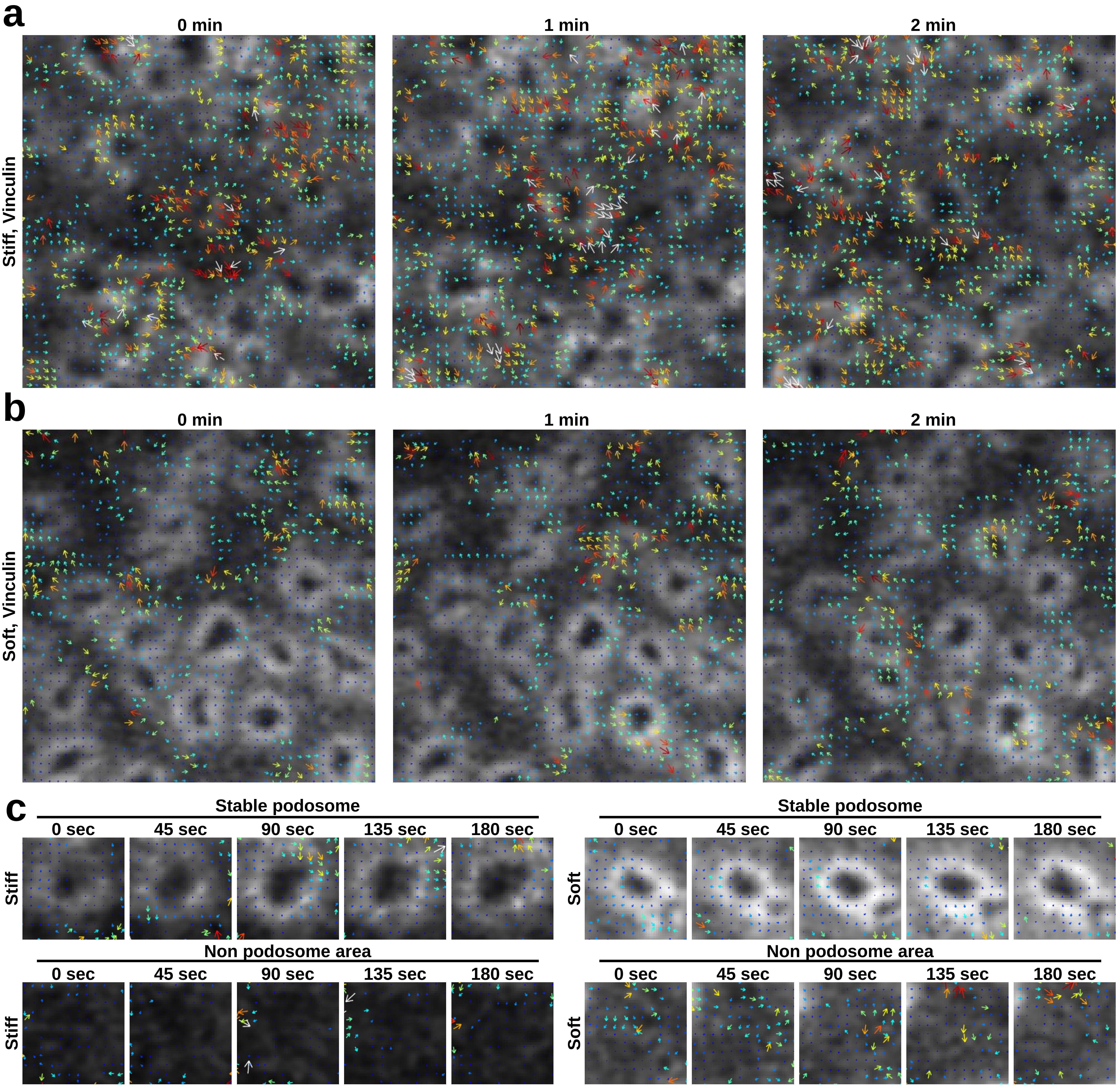


**Supplementary Figure 14**

**twSTICS reveals flows of vinculin in dynamic podosomes on stiff and soft substrates**

**a-c,** DCs were transfected with Lifeact-GFP and vinculin-mCherry. Airyscan imaging was performed with 15s frame intervals. Time series were subjected to twSTICS analysis and results are plotted as vector maps in which the arrows indicate direction of flow and both the size and color are representative of the flow magnitude. Shown are representative waves of vectors for vinculin on **a,** stiff and **b,** soft substrates as well as in **c,** stable and non podosome areas.

### Van den Dries et al. Supplementary Figure 15

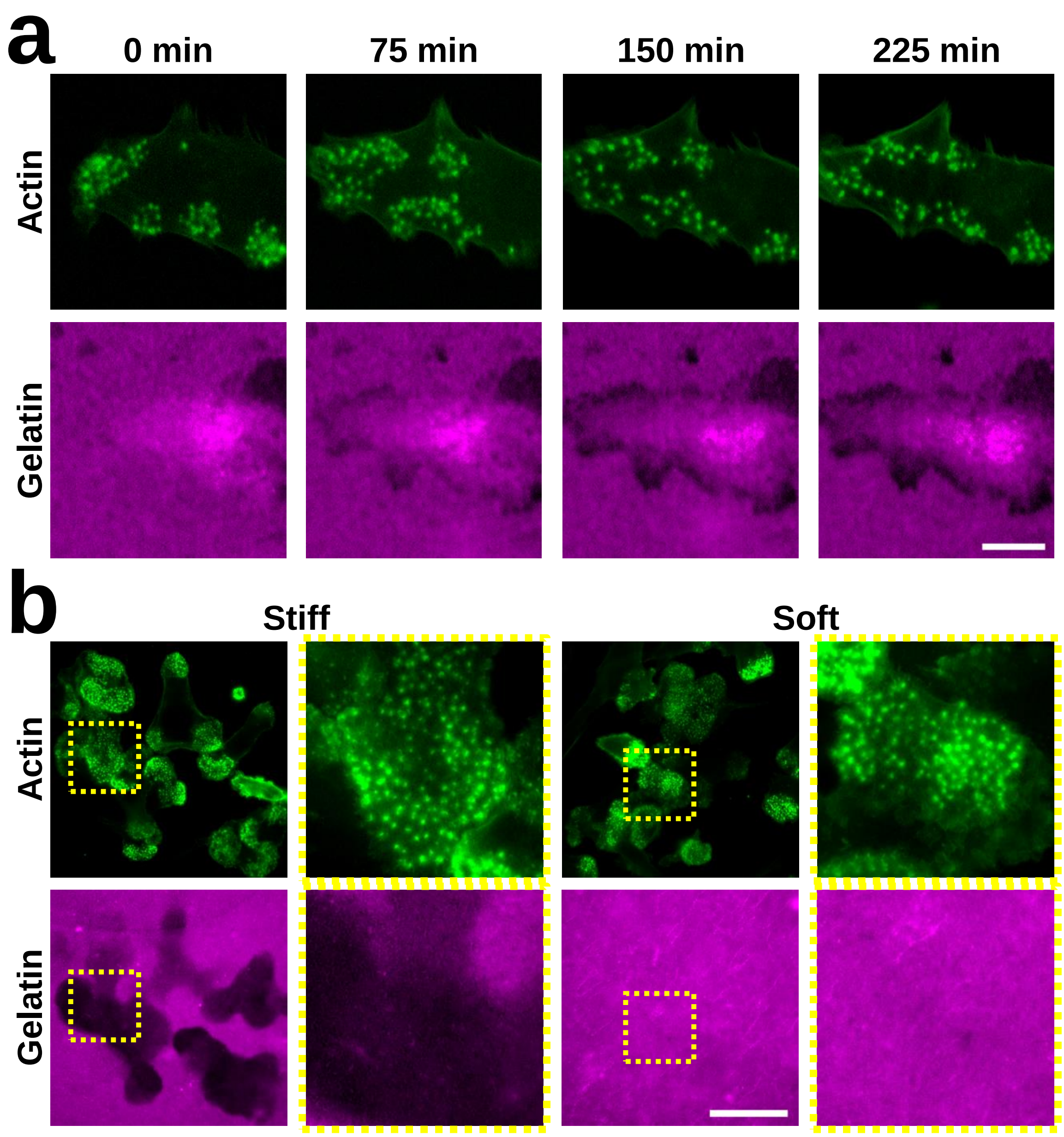


**Supplementary Figure 15**

**Decreased gelatin degradation on soft substrates**

**a**, DCs were transfected with Lifeact-GFP (green) and seeded on stiff PDMS that as coated with rhodamine-conjugated gelatin (magenta). Image series were acquired overnight on a Leica DM6000 widefield microscope with 1 image per 5 min. Shown are stills from the image series demonstrating the specific degradation of gelatin underneath areas with podosome formation (See also Supplementary Video 6). **b**, DCs were seeded on gelatin-rhodamine (magenta) coated stiff and soft substrates, incubated overnight and subsequently stained for actin (green). Shown are representative images of gelatin degradation of stiff (right panels) and soft (left panels) substrates. Images in column 1 and 3 are the same as in Fig. 5j. Images in column 2 and 4 depict representative single cells. Scale bars: **a** = 10 µm, **b** = 20 µm.

### Van den Dries et al. Supplementary Figure 16

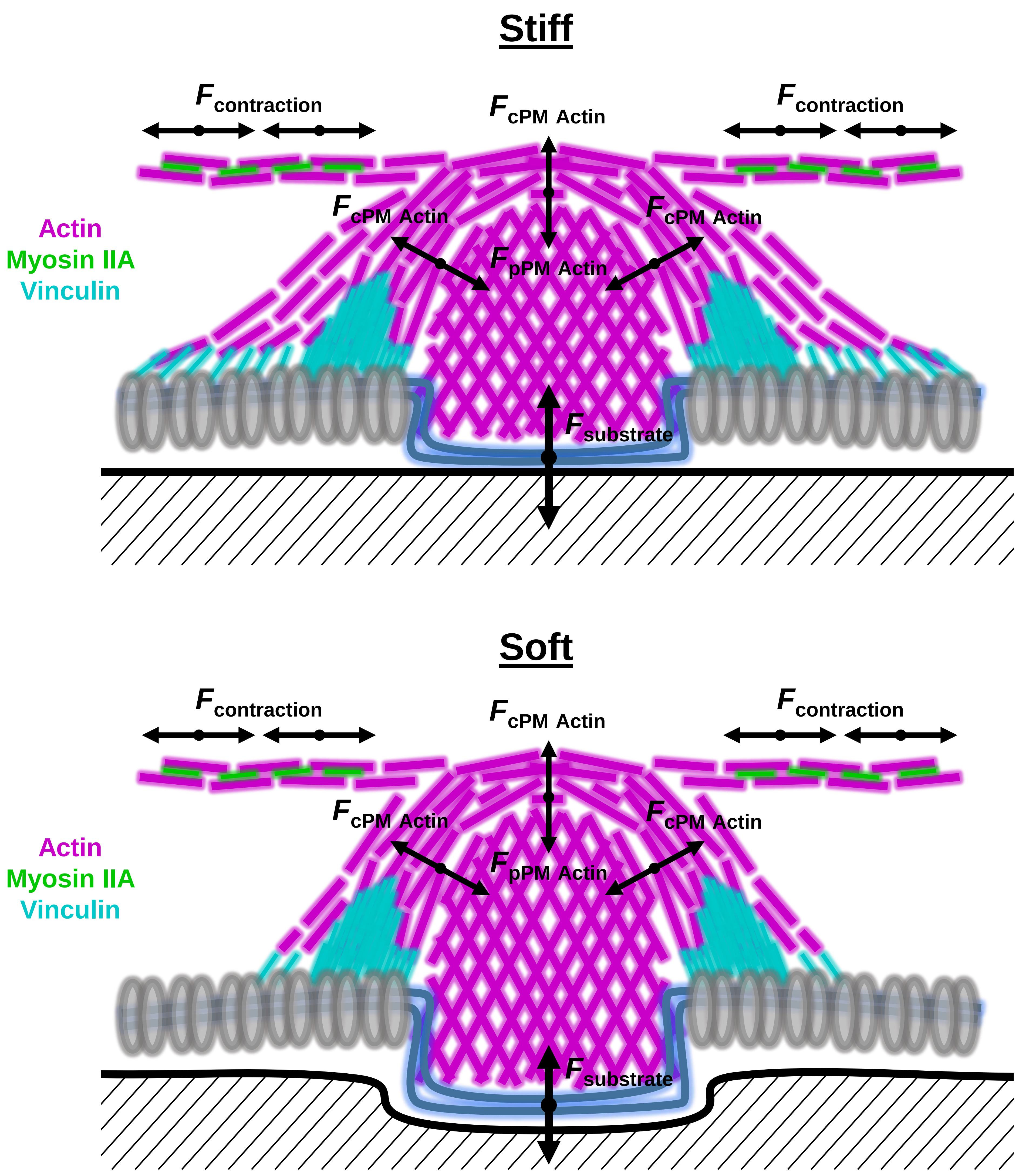


**Supplementary Figure 16**

**Schematic model of how forces may be balanced in the podosome protrusive core**
cPM actin polymerization generates a downward protrusive force that is initially counterbalanced by the an upward force from the underlying substrate. The subsequent vertical growth of the cPM actin generates an upward force at the top of podosomes that is balanced by the pPM actin. Myosin IIA in the dorsal actin filaments induces contractile forces, probably to facilitate long range inter-podosomal force connections.

### Van den Dries et al. Supplementary Figure 17

a

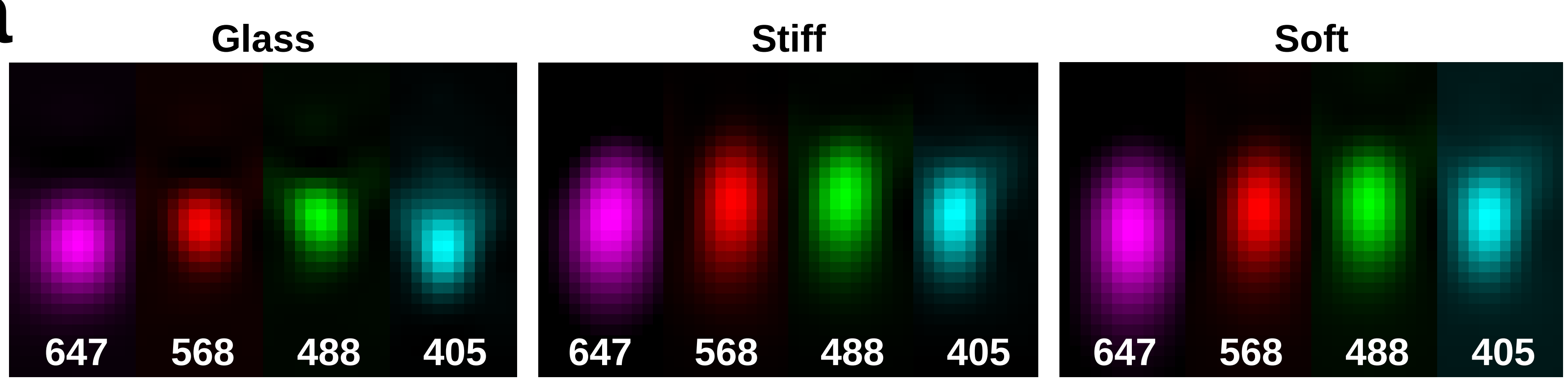


b

Glass

| | FWHM (XY) | FWHM (Z) |
|---|---|---|
| **405** | 185 nm | 611 nm |
| **488** | 174 nm | 511 nm |
| **568** | 194 nm | 522 nm |
| **647** | 265 nm | 705 nm |

Stiff

| | FWHM (XY) | FWHM (Z) |
|---|---|---|
| **405** | 190 nm | 740 nm |
| **488** | 197 nm | 818 nm |
| **568** | 199 nm | 910 nm |
| **647** | 261 nm | 1070 nm |

Soft

| | FWHM (XY) | FWHM (Z) |
|---|---|---|
| **405** | 188 nm | 741 nm |
| **488** | 194 nm | 818 nm |
| **568** | 198 nm | 861 nm |
| **647** | 250 nm | 1080 nm |

**Supplementary Figure 17**

**PSF measurements on glass, stiff and soft PDMS**

**a,** TetraSpeck microspheres of 200 nm labeled with Alexa405, Alexa488, Alexa568 and Alexa647 were left to settle down on glass, stiff and soft PDMS and PSF measurements were collected using Airyscan settings on the LSM 880. Shown are the x-y,z profiles of the beads for each of the four channels. **b,** Quantification of the FWHM values in x-y as well as in z for the four channels on glass, stiff and soft PDMS. Note that the FWHM is not altered by PDMS in x-y but is increased by approximately 1.5 fold in z compared to glass.

#### Supplementary Video Legends

##### Supplementary Video 1

Assembling podosome in a DC transfected for cortactin-BFP, α-actinin-tagRFP, vinculin-GFP and Lifeact-iRFP. Image series were acquired on a Leica DMI6000 with 1 image per 6 s. Playback speed: 15 fps

##### Supplementary Video 2

DCs were transfected with Lifeact-GFP and vinculin-mCherry and seeded on stiff substrates. Image series were acquired on a Zeiss LSM880 with Airyscan settings with 1 image per 15 s. Playback speed: 15 fps

##### Supplementary Video 3

DCs were transfected with Lifeact-GFP and vinculin-mCherry and seeded on soft substrates. Image series were acquired on a Zeiss LSM880 with Airyscan settings with 1 image per 15 s. Playback speed: 15 fps

##### Supplementary Video 4

twSTICS reveals vinculin and actin flows in podosome clusters in a DC seeded on stiff PDMS.

##### Supplementary Video 5

twSTICS reveals vinculin and actin flows in podosome clusters in a DC seeded on soft PDMS.

##### Supplementary Video 6

DCs were transfected with Lifeact-GFP and seeded on rhodamine-gelatin labeled stiff substrates. Image series were acquired on a Leica DMI6000 with 1 image per 5 min. Playback speed: 15 fps
